# Isolation and Diversity of Sediment Bacteria in the Hypersaline Aiding Lake, China

**DOI:** 10.1101/638304

**Authors:** Tong-Wei Guan, Yi-Jin Lin, Meng-Ying Ou, Ke-Bao Chen

## Abstract

A total of 343 bacteria from sediment samples of Aiding Lake, China, were isolated using nine different media with 5% or 15% (w/v) NaCl. The number of species and genera of bacteria recovered from the different media significantly varied, indicating the need to optimize the isolation conditions. The results showed an unexpected level of bacterial diversity, with four phyla (Firmicutes, Actinobacteria, Proteobacteria, and Rhodothermaeota), fourteen orders (Actinopolysporales, Alteromonadales, Bacillales, Balneolales, Chromatiales, Glycomycetales, Jiangellales, Micrococcales, Micromonosporales, Oceanospirillales, Pseudonocardiales, Rhizobiales, Streptomycetales, and Streptosporangiales), including 17 families, 41 genera, and 71 species. In this study, the predominant phyla included Firmicutes and Actinobacteria and the predominant genus included *Halomonas*, *Gracilibacillus*, *Streptomyces*, and *Actinopolyspora*. To our knowledge, this is the first time that members of phylum Rhodothermaeota were identified in sediment samples from a salt lake. This study has identified at least four novel isolates.

## 1 Introduction

Halophiles thrive in hypersaline niches and have potential applications in biotechnology[1, 2] (Oren 2010, 2015). Microbial diversity in most hypersaline environments is often studied using culture-dependent and -independent methods[3–7]. Previous studies have shown that the taxonomic diversity of microbial populations in terrestrial saline and hypersaline environments is relatively low [8, 9]. Halophilic microbial communities vary with season [10], and in general, microbial diversity decreases with increased salinity [11, 12]. Hypersaline lakes are considered extreme environments for microbial life. A variety of salt lakes have been surveyed for bacterial diversity such as Chaka Lake in China, Chott El Jerid Lake in Tunisia, Meyghan Lake in Iran, Keke Lake in China, and Great Salt Lake in the United States [5, 13–16]. In addition, groups of novel halophilic or halotolerant bacteria in salt lakes have been described using culture-dependent methods such as *Brevibacterium salitolerans* sp. nov., *Amycolatopsis halophila* sp. nov., *Actinopolyspora lacussalsi* sp. nov., *Halomonas xiaochaidanensis* sp. nov., *Salibacterium nitratireducens* sp. Nov., and *Paracoccus halotolerans* sp. nov. [17–22]. Despite these previous studies, our understanding of bacterial diversity in hypersaline lakes remains limited, particularly in athalassohaline lakes at low elevations. Aiding Lake represents an ideal site for studying halophilic or halotolerant bacteria in a hypersaline lake. The salt lake is located on the Turpan Basin (the hottest place in China) and possesses a salinity of 31.6%. In fact, our current understanding of the bacterial diversity in the Aiding Lake using a culture-dependent method is limited. To our knowledge, this is the first attempt to comprehensively characterize the bacteria diversity in dry salt lake sediments. The aim of the study was to investigate the bacterial diversity and to mine novel bacterial species from Aiding Lake, China.

## 2 Materials and methods

### 2.1 Site description and sample collection

Aiding Lake is a dry salt lake located on the Turpan Basin in northwestern China, with an elevation of 154 m–293 m below sea level. The high salinity, low nutrient levels, dry climate, and high UV intensity makes it an extreme environment. Aiding Lake covers an area of about 60 km^2^ and is a closed ecosystem, without the influx of perennial rivers. Three soil samples, namely, S1 (89°21’98”E, 42°40’42”N), S2 (89°16’6”E, 42°38’55”N), and S3 (89°20’26”E, 42°41’53”N) were collected from the lake sediments. Soil sample temperature is 23.6°C-25.1°C and the pH is approximately 7.5–8.6. Three sediments samples were collected from a depth of 1 to 30 cm in mid-July of 2012. The distance between two sample points was greater than 5 km. Samples were stored at 4°C in the field and immediately transported to the laboratory. The concentrations of major cations and trace elements in the dissolved sediments were measured according to Yakimov et al. (2002) [23].

### 2.2 Isolation of microorganisms

Three sediment samples were selected for cultivation of bacteria. To isolate halophilic and/or halotolerant bacteria, the sediments (10 g wet weight) were dispersed into 90 mL of sterilized NaCl brine (5% or 15%, w/v) and incubated at 37°C for 60 min with shaking at 200 rpm. The resulting slurry was then serially diluted with sterilized NaCl brine (5% or 15%, w/v). Aliquots (0.1 mL) of each dilution were spread onto Petri dishes using nine media (Table 1). The compositions of the nine media used for the isolation of bacteria from the Aiding Lake samples are shown in Table 1. All agar plates were supplemented with 5% or 15% (w/v) NaCl. To suppress the growth of nonbacterial fungi, the solidified media were supplemented with nystatin (50 mg·L^−1^). The Petri dishes were incubated at 37°C for one to six weeks. Based on size and color, colonies were picked and further purified on inorganic salts-starch agar [24] or TSA supplemented with 5% or 15% (w/v) NaCl, and as glycerol suspension (20%, v/v) at −20 °C or as lyophilized cells for long-term storage at −4 °C.

**Table 1.** Compositions of the nine different media used for the isolation of bacteria from Aiding Lake samples.

| Medium | Composition | Reference |
| --- | --- | --- |
| A | Inorganic salts-starch agar (ISP 4) | Shirling and Gottlieb 1966 |
| B | Casein hydrolysate acid-starch agar: starch 5.0 g, casein hydrolysate acid 0.5 g, KNO<sub>3</sub> 0.5 g, Aspartic acid 0.1g, CaCO<sub>3</sub>, 0.3 g, K<sub>2</sub>HPO<sub>4</sub> 0.5 g, MgCl<sub>2</sub> 0.2 g, FeSO<sub>4</sub>·7H<sub>2</sub>O 10 mg, agar 18 g | This study |
| C | Microcrystalline cellulose-proline agar: microcrystalline cellulose 2.0 g, proline 0.5 g, arginine 0.1 g, KCl 10 g, (NH<sub>4</sub>)<sub>2</sub>SO<sub>4</sub> 1.0 g, K<sub>2</sub>HPO<sub>4</sub> 0.2 g, CaCO<sub>3</sub> 0.02 g, MgSO<sub>4</sub> · 7H<sub>2</sub>O 2 g, FeSO<sub>4</sub> ·7H<sub>2</sub>O 10 mg, MnCl<sub>2</sub>·4H<sub>2</sub>O 1mg, agar 18 g | This study |
| D | Glycerin-asparagine agar: glycerin 3g, asparagine 1g, C<sub>3</sub>H<sub>3</sub>NaO<sub>3</sub> 0.5g, MgSO<sub>4</sub>· 7H<sub>2</sub>O 2 g, FeSO<sub>4</sub>·7H<sub>2</sub>O 10 mg, ZnSO<sub>4</sub>·7H<sub>2</sub>O 1 mg, VB1 0.1 mg,VB6 0.05 mg, biotin 0.2mg, agar 18 g | This study |
| E | Yeast extract-casamino acids agar: yeast extract 3g, Casamino acids 2g, Sodium glutamate 1g, Trisodium citrate 1g, MgSO<sub>4</sub>·7H<sub>2</sub>O 5g, CaCl<sub>2</sub>·2H<sub>2</sub>O 1g, KCl 3g, FeCl<sub>2</sub>·4H<sub>2</sub>O 0.2mg, MnCl<sub>2</sub>·4H<sub>2</sub>O 0.2mg, agar, 18 g | This study |
| F | Stachyose tetrahydrate-Alanine agar: stachyose tetrahydrate 5g, alanine 2g, KNO<sub>3</sub> 0.2g, CaCO<sub>3</sub> 0.02g, MgSO<sub>4</sub>·7H<sub>2</sub>O 0.05g, KCl 20g, FeSO<sub>4</sub>·7H<sub>2</sub>O 10 mg, MgCl<sub>2</sub> 20g, agar 18 g | This study |
| G | Microcrystalline cellulose-sorbitol agar: microcrystalline cellulose 10g, Sorbitol 2g, Beta-Cyclodextrin 1g, MgSO<sub>4</sub>·7H<sub>2</sub>O 0.1g , CaCO<sub>3</sub> 0.5g , FeSO<sub>4</sub> 0.01g , KCl 20g, MgCl<sub>2</sub> 10g, agar 18 g | This study |
| H | Yeast extract-fish peptone agar: yeast extract 1g, fish peptone 0.5g, NH<sub>4</sub>Cl 0.5g, MgSO<sub>4</sub>·7H<sub>2</sub>O 20g, MgCl<sub>2</sub>·6H<sub>2</sub>O 15g, KCl 5g, sodium pyruvate 1g, K<sub>2</sub>HPO<sub>4</sub> 0.3g, CaCl<sub>2</sub>·2H<sub>2</sub>O 0.2g, agar 18 g | This study |
| I | Yeast extract-glycerin agar: yeast extract 10g, Glycerin 0.5g, peptone 0.5g, (NH<sub>4</sub>)NO<sub>3</sub> 0.1g, MgCl<sub>2</sub> 5g, Na<sub>2</sub>SO<sub>4</sub> 3g, yeast KCl 1g, NaHCO<sub>3</sub> 2g, KBr 0.05g, SrCl<sub>2</sub> 0.01g, Na<sub>2</sub>SiO<sub>3</sub> 0.001g, agar 18 g | This study |

### 2.3 Identification of bacteria

Isolates were tentatively grouped and dereplicated by observing their morphological and culture characteristics, including colony features (color, shape, and size) on plates and slants, the presence of aerial mycelia and substrate mycelia, distinctive reverse colony color, diffusible pigment, salinity tolerance (5% or 15% NaCl, w/v), spore mass color, and sporophore and spore chain morphology. Based on the preliminary grouping, 71 isolates were selected and subjected to 16S rRNA gene sequence analysis for precise genus and species identification. The identities of the organisms were determined based on nearly full-length 16S rRNA gene sequence analysis. Genomic DNA was extracted from each isolate, and the 16S rRNA gene sequence was amplified as described by Li et al. (2007) [25] with primers PA (5’-CAGAGTTTGATCCTGGCT-3’) and PB (5’-AGG AGGTGATCCAGCCGC A-3’), or the primers 27F (5’-AGAGTTTGATCMTGGCTCAG-3’) and 1492R (5’-GGTTACCTTGTT ACGACTT-3’). PCR products were purified using a PCR purification kit (Sangon, Shanghai, China). The almost-complete 16S rRNA gene sequence of 71 strains was obtained. Multiple alignments with sequences of the most closely related recognized species and calculations of levels of sequence similarity were conducted using EzBioCloud server [26]. Phylogenetic analysis was performed using the software package MEGA version 6.0 [27]. Phylogenetic trees were constructed according to the neighbor-joining method [28]. Evolutionary distance matrices were generated as described by Kimura (1980) [29]. The topology of the phylogenetic tree was evaluated using the bootstrap resampling method of Felsenstein (1985) [30] with 1000 replicates. DNA-DNA relatedness values were determined using the fluorometric microwell method [31].

### 2.4 Nucleotide sequence accession numbers

The sequences of the bacterial isolates reported in this study have been deposited to GenBank (Accession no. MK818765-MK818834, MK296404).

## 3 Results

### 3.1 Sediment geochemistry

Physicochemical parameters were distinct among the three sediment samples (Table 2). In the S1, S2, and S3 samples, Na^+^ concentrations ranged from 26.37 g/Kg to 83.93 g/Kg and Cl^−^ concentrations ranged from 33.28 g/Kg to 389.11 g/Kg, which are typical of chloride-type environments. The pH of the sediment samples ranged from 7.6 to 8.3, indicating a slightly alkaline environment. In sample S3, ionic composition (e.g., Mg^2+^, Na^+^, K^+^, Mn^2+^, and Cl^−^) was significantly lower than the other samples. The other physicochemical properties of the samples are presented in Table 2.

**Table 2.**
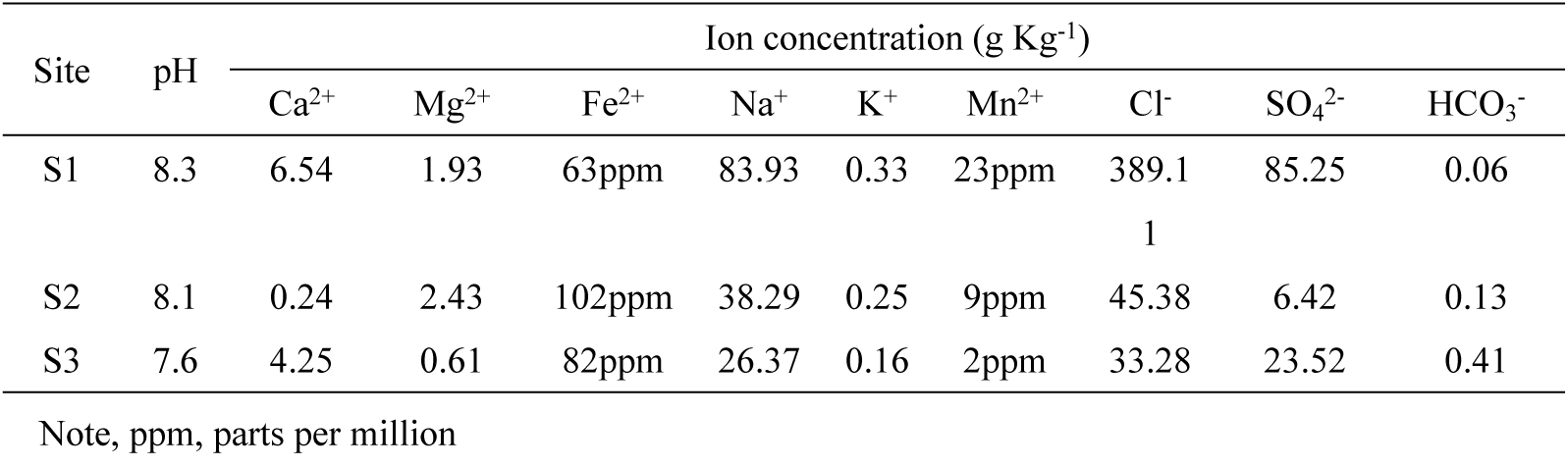
Physicochemical properties of the sediments from the three sample sites in Aiding Lake.

### 3.2 Phylogenetic analysis of bacterial isolates

A total of 343 strains were isolated using different media. The 16S rRNA gene of 71 chosen isolates was sequenced. The percentages of 16S rRNA gene sequence similarities (91.14% to 100%) of these isolates to the closest type strains are presented in Table 3. The bacteria isolated in this study displayed considerable diversity. The predominant phyla were Firmicutes (149 strains, 43.4%) and Actinobacteria (121 strains, 35.3%). The other bacterial isolates belonged to phyla Proteobacteria (68 strains, 19.8%) and Rhodothermaeota (5 strains, 1.5%). The isolates were distributed among 14 orders, namely, Actinopolysporales (26 strains), Alteromonadales(8 strains), Bacillaceae (149 strains), Balneolales (5 strain), Chromatiales (12 strain), Glycomycetales (2 strains), Jiangellales (3 strains), Micrococcales (8 strain), Micromonosporineae (5 strains), Oceanospirillales (45 strains), Pseudonocardiales (11 strains), Rhizobiales (3 strain), Streptomycetales (27 strains), and Streptosporangiales (26 strain), including 41 genera and 71 species (Tables 3 and 4). Other organisms (ADL013 and ADL023) could not be accurately identified to the genus level. In the study, *Halomonas* (42 strains, 12.3%), *Gracilibacillus* (33 strains, 9.7%), *Streptomyces* (27 strains, 7.9%), *Actinopolyspora* (26 strains, 7.6%), *Nocardiopsis* (17 strains, 4.9%), *Aquisalimonas* (12 strains, 3.5%), *Saccharomonospora* (12 strains, 3.5%), *Bacillus* (12 strains, 3.5%), *Marinococcus* (11 strains, 3.2%), *Virgibacillus* (11 strains, 3.2%), and *Sediminibacillus* (11 strains, 3.2%) were some dominant group in sediment samples of Aiding Lake, the number of microorganisms in each of the other genus is relatively small (Table 3 and Fig. 1). For example, *Anaerobacillu*, *Ornithinibacillus*, and *Prauserella* include only one strain, respectively. To isolate halophilic or halotolerant bacteria in the sediments of Aiding Lake, the agar plates were supplemented with 5% or 15% (w/v) NaCl. Approximately 141 strains isolated from these media with 5% NaCl belonged to 25 different genera, and 202 strains isolated from the media using 15% NaCl belonged to 23 different genera (Table 3).

**Table 3.**
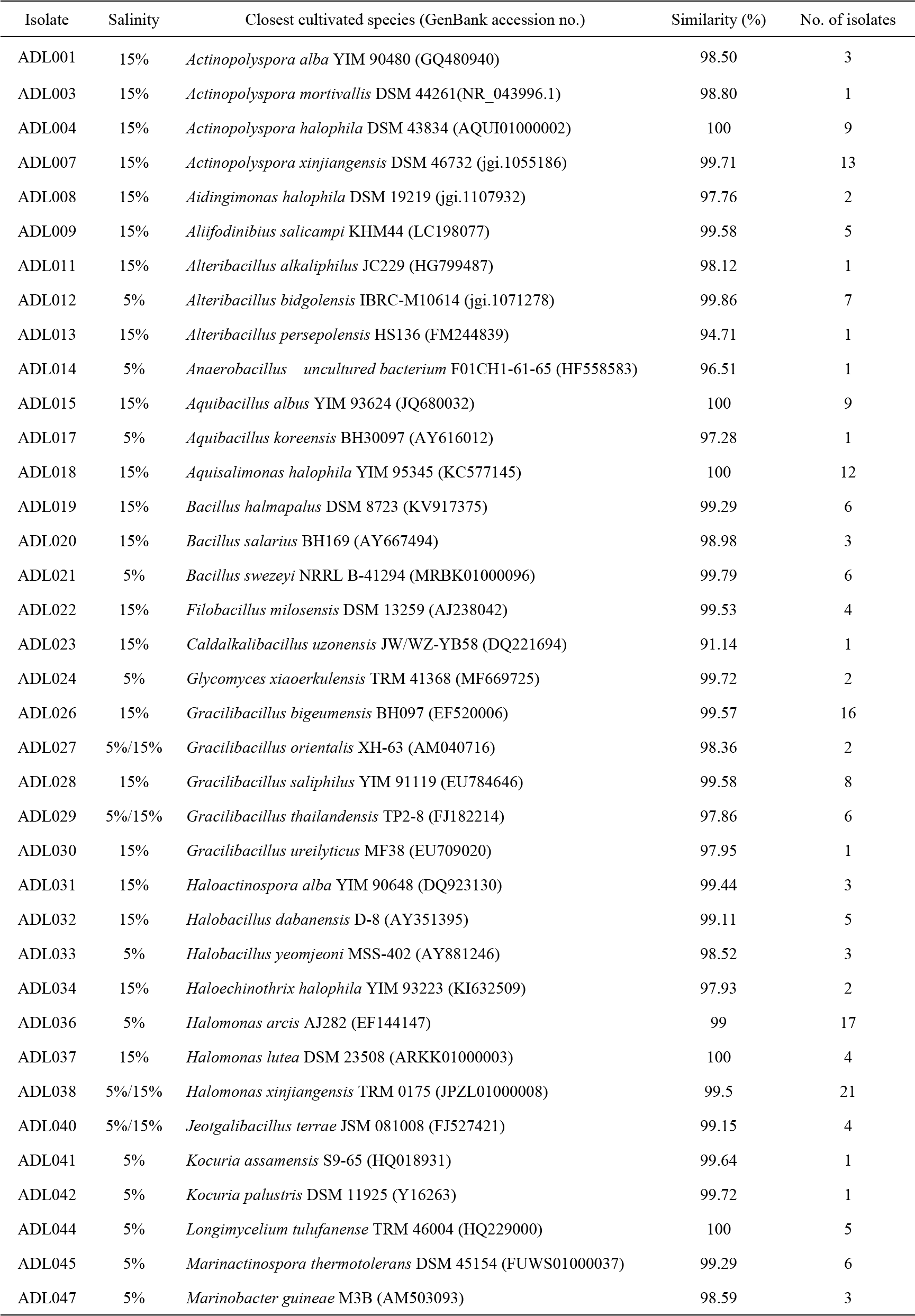
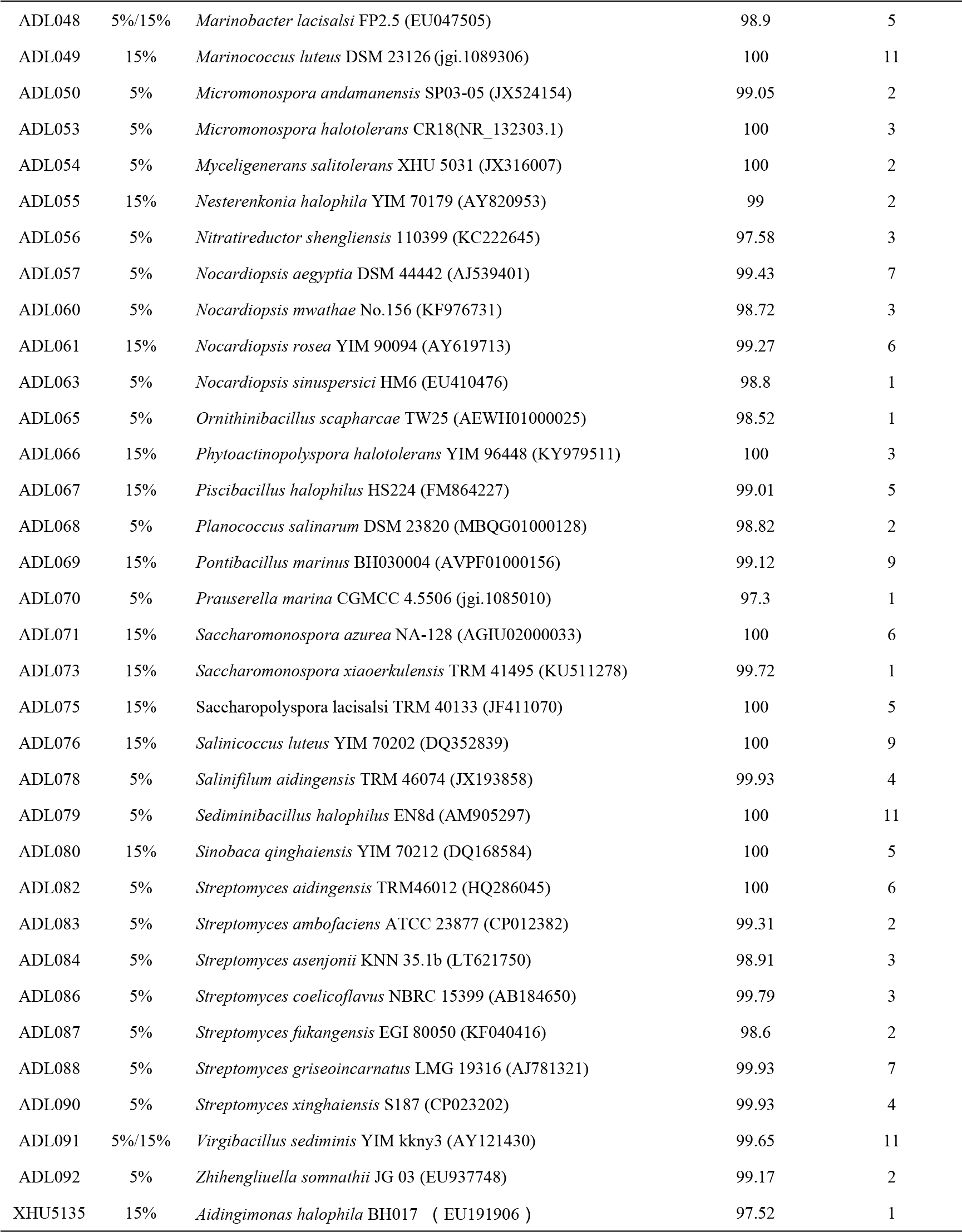
Bacteria isolated using two different salinity from sediments of Aiding Lake, with the similarity values for 16S rRNA gene sequences.

**Table 4.**
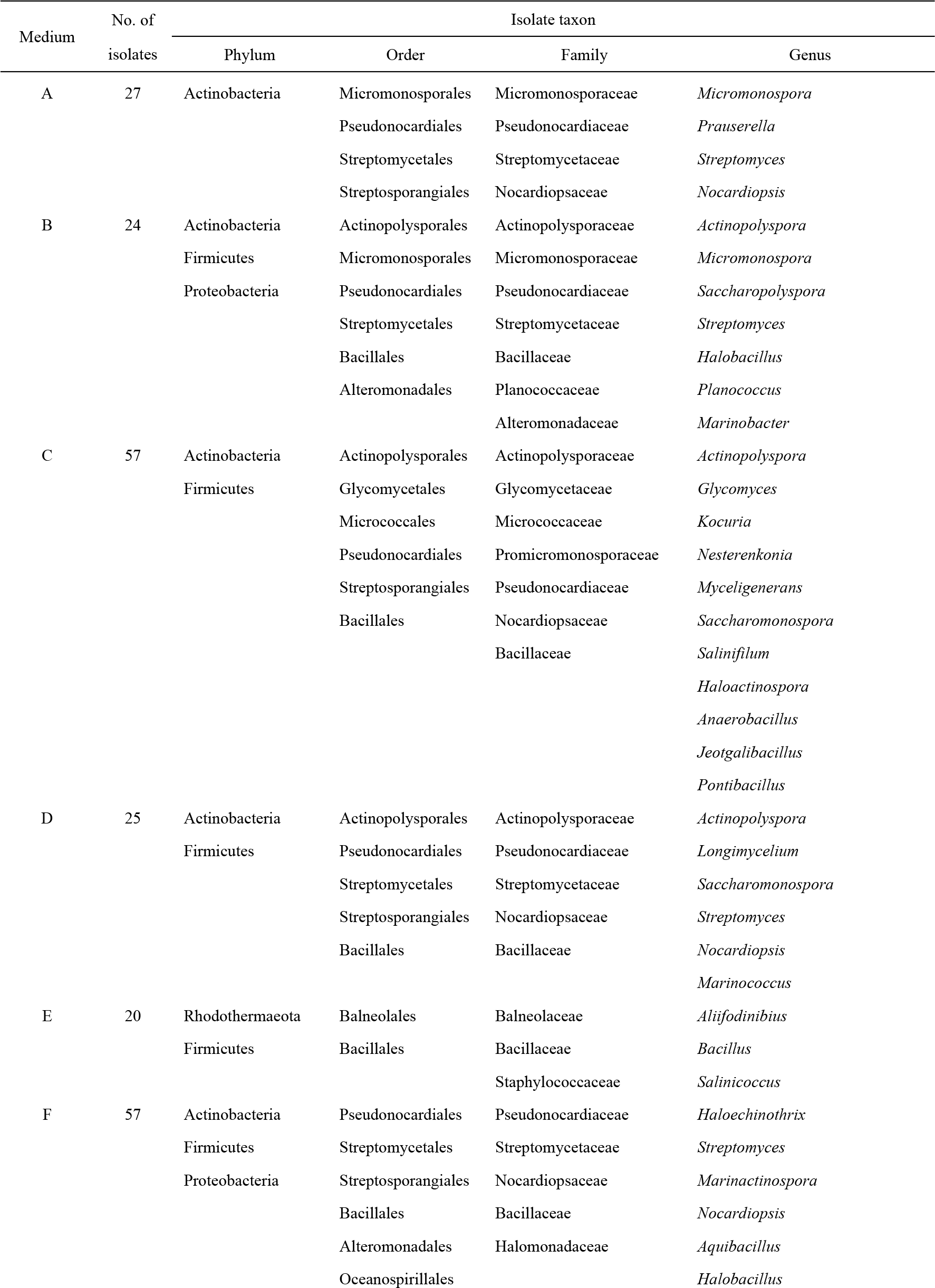
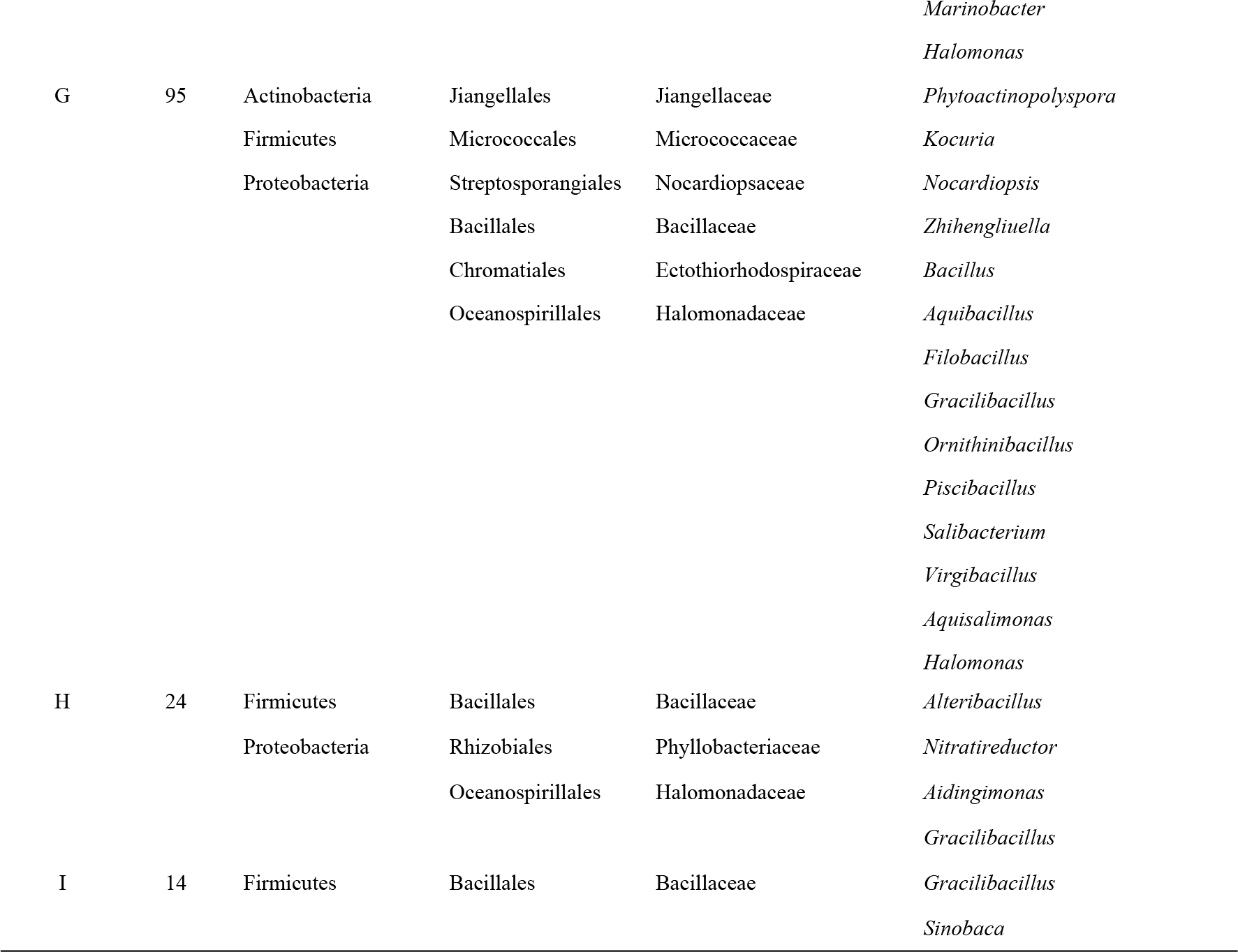
Statistical analyses of the relationships between the taxa of the bacterial strains and the nine different media.

**Fig 1.**
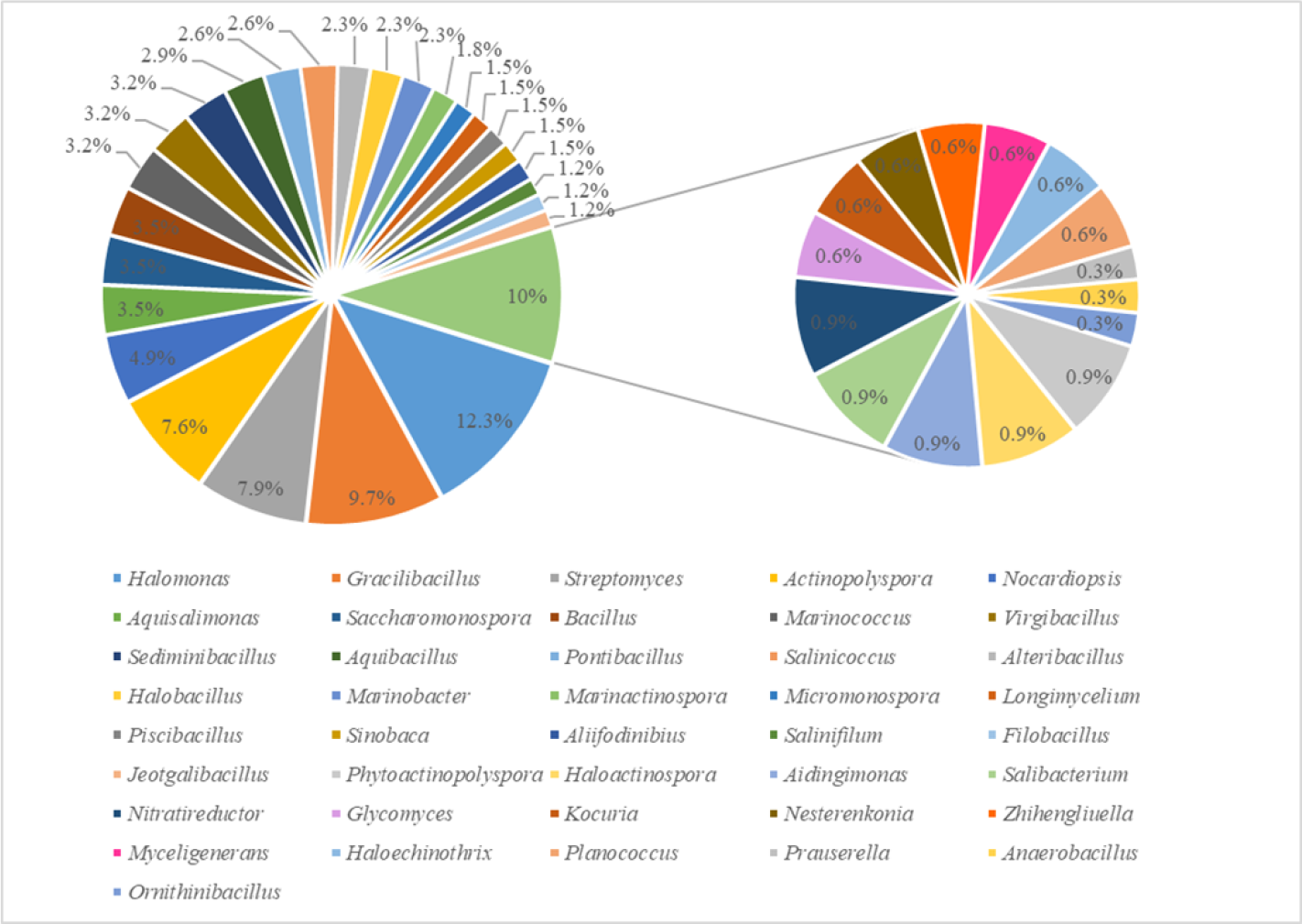
Percentage of isolated strains in each genus.

### 3.3 Bacterial isolates from different media

To obtain additional bacterial groups, the sediment samples from Aiding Lake were isolated using nine different media (Table 1). Five class of bacteria, namely, Actinobacteria, Bacilli, Alphaproteobacteria, Gammaproteobacteria, and Balneolia, including 14 orders, 17 family and 41 genera, were obtained (Table 4). Most of the bacterial groups were isolated using microcrystalline cellulose-sorbitol agar (G), and 14 bacterial genera (*Aquibacillus*, *Aquisalimonas*, *Bacillus*, *Filobacillus*, *Gracilibacillus*, *Halomonas*, *Kocuria*, *Nocardiopsis*, *Ornithinibacillus*, *Phytoactinopolyspora*, *Piscibacillus*, *Salibacterium*, *Virgibacillus*, and *Zhihengliuella*) were isolated. At the same time, microcrystalline cellulose-proline agar (C), stachyose tetrahydrate-alanine agar (F), casein hydrolysate acid-starch agar (B), and glycerin-asparagine agar (D) resulted in relatively efficient isolations for 11 genera (*Actinopolyspora*, *Glycomyces*, *Kocuria*, *Nesterenkonia*, *Myceligenerans*, *Saccharomonospora*, *Salinifilum*, *Haloactinospora*, *Anaerobacillus*, *Jeotgalibacillus*, and *Pontibacillus*), 8 genera (*Haloechinothrix*, *Streptomyces*, *Marinactinospora*, *Nocardiopsis*, *Aquibacillus*, *Halobacillus*, *Marinobacter*, and *Halomonas*), 7 genera (*Actinopolyspora*, *Micromonospora*, *Saccharopolyspora*, *Streptomyces*, *Halobacillus*, *Planococcus*, and *Marinobacter*), and 6 genera (*Actinopolyspora*, *Longimycelium*, *Saccharomonospora*, *Streptomyces*, *Nocardiopsis*, and *Marinococcus*), respectively. Only two genera (*Gracilibacillus* and *Sinobaca*) were isolated using yeast extract-glycerin agar (I). The results of isolation using the other media are shown in Table 4. From the number of isolated strains, 95, 57, and 57 strains were isolated using media G, F, and C, respectively, and the number of strains isolated from media A, B, D, E, H and I was relatively small (Table 4). In addition, the bacterial diversity recovered using different media also varied considerably. Medium G had the best recoverability, with 14 of the total of 42 bacterial genera recovered. Although media I and E showed the lowest recoverability at the genus level (Table 4), the microorganism of *Sinobaca* genus was only isolated using medium I, and a novel strain ADL013 was found using the medium too. At the same time, the microorganism of phylum Rhodothermaeota was detected only using medium E (Table 3). I n short, to obtain more bacterial resources, it is also necessary to develop different types of media.

## 4 Discussion

We isolated and cultured 343 representatives of 41 bacterial genera. To our knowledge, this is the first study to recover such a high diversity of culturable bacteria from salt lake sediments. Bacterial diversity in Aiding Lake was based on the 16S rRNA gene sequences, which was relatively higher than other salt lakes at the genus level. For example, sequencing of 16S DNA indicated the presence of members of bacterial genera *Bacillus*, *Halomonas*, *Pseudomonas*, *Exiguobacterium*, *Vibrio*, *Paenibacillus*, and *Planococcus* in the salt lake La Sal del Rey, in extreme South Texas (USA) [32]. There are 152 strains affiliated with the 11 genera: *Bacillus*, *Gracilibacillus*, *Halobacillus*, *Halolactibacillus*, *Thalassobacillus*, *Virgibacillus*, *Salimicrobium*, *Staphylococcus*, *Halomonas*, *Idiomarina*, and *Chromohalobacter* from Yuncheng Salt Lake, China [33]. Nine bacterial genera (*Salicola*, *Halovibrio*, *Halomonas*, *Oceanobacillus*, *Thalassobacillus*, *Halobacillus*, *Virgibacillus*, *Gracilibacillus*, *Salinicoccus*, and *Piscibacillus*) were isolated from Howz Soltan Lake, Iran [34]. Notably, some rare genera from the sediments of Aiding Lake such as *Aliifodinibius*, *Aidingimonas*, *Filobacillus*, *Haloechinothrix*, *Jeotgalibacillus*, *Longimycelium*, *Myceligenerans*, *Ornithinibacillus*, *Phytoactinopolyspora*, and *Piscibacillus* were discovered in the present study. In addition, phylum Rhodothermaeota was detected for the first time in sediment samples from a salt lake.

The richness and diversity of actinobacteria isolated from Aiding Lake in the present study were relatively high, consiting of eight orders (Actinopolysporales, Glycomycetales, Jiangellales, Micrococcales, Micromonosporales, Pseudonocardiales, Streptomycetales and Streptosporangiales), including 18 genera: *Actinopolyspora*, *Glycomyces*, *Haloactinospora*, *Haloechinothrix*, *Kocuria*, *Longimycelium*, *Marinactinospora*, *Micromonospora*, *Myceligenerans*, *Nesterenkonia*, *Nocardiopsis*, *Prauserella*, *Phytoactinopolyspora*, *Saccharomonospora*, *Saccharopolyspora*, *Salinifilum*, *Streptomyces*, and *Zhihengliuella*. Our results indicated that the diversity of bacterial 16S rRNA gene sequences of strains from Aiding Lake were more diverse at the genus level than those reported in saline environment. Fourteen genera, namely *Micromonospora*, *Nocardia*, *Streptomyces*, *Nocardiopsis*, *Saccharopolyspora*, *Pseudonocardia*, *Verrucosispora*, *Mycobaterium*, *Actinomuaura*, *Actinomycetospora*, *Streptosprangium*, *Microbispora*, and *Sphaerosporangium* were isolated from marine sponges in Florida, USA [35]. Ten genera, including *Actinotalea*, *Arthrobacter*, *Brachybacterium*, *Brevibacterium*, *Kocuria*, *Kytococcus*, *Microbacterium*, *Micrococcus*, *Mycobacterium*, and *Pseudonocardia* from Arctic marine sediments were obtained by culture-dependent approaches [36]. Fifty-two halophilic actinomycetes belonging to the *Actinopolyspora*, *Nocardiopsis*, *Saccharomonospora*, *Streptomonospora*, and *Saccharopolyspora* genera, were isolated from Saharan soils of Algeria [37]. The genera *Dietzia*, *Microbacterium*, *Rhodococcus*, and *Nocardia* of actinobacteria were isolated from Lake Magadi, Kenya [38].

Among 343 strains isolated, the 16S rRNA gene sequences of 71 isolated halophilic or halotolerant bacteria were compared to those deposited in the public database (EzBioCloud, https://www.ezbiocloud.net/identify). As the results indicated, most of the strains exhibited > 97% similarity to other published type species (Table 3). However, strain ADL014 shared 96.51% similarity with *Anaerobacillus alkalidiazotrophicus* F01CH1-61-65. It is generally accepted that organisms displaying 16S rRNA sequence similarity values of 97% or less belong to different species [39]. Sequence analysis indicated that strain ADL014 formed a distinct lineage within the genus *Anaerobacillus* and always had the closest phylogenetic affinity to members of the genus *Anaerobacillus* (Fig 2). Phylogenetic reconstruction indicated that strain ADL014 represents a novel species. Strain ADL013 exhibited 94.71% similarity to the 16S rRNA gene sequence of *Alteribacillus persepolensis* HS136. Phylogenetic analysis also showed that strain ADL013 can be distinguished from representatives of genera in the family Bacillaceae, and strain ADL013 formed a distinct lineage within family Bacillaceae (Fig 3). The strain had such low degrees of sequence similarity, suggesting these might be a novel species or genus of Bacillaceae. Strain ADL023 exhibited 91.14% similarity to the 16S rRNA gene sequence of *Caldalkalibacillus uzonensis* JW/WZ-YB58. Phylogenetic analysis showed that strain ADL023 formed a distinct lineage within family Bacillaceae (Fig 4), and can be well distinguished from representatives of genera in family Bacillaceae. Therefore, it could represent a new genus in family Bacillaceae. The 16S rRNA gene sequences of XHU5135 showed 97.52% identities to the nearest neighbors, *Aidingimonas halophila* YIM 90637 [40]. Although it showed higher 16S rRNA gene similarities (97.52%) to the closest recognized strains, DNA-DNA hybridization experiments revealed that levels of DNA-DNA relatedness between strain XHU 5135 and *Aidingimonas halophila* YIM 90637 were 23.3± 2.8%. This values is below the 70% cut-off point recommended for recognition of genomic species [41]. Thus, strain XHU 5135 represents a novel species of the genus *Aidingimonas*. The neighbour-joining phylogenetic tree based on 16S rRNA gene sequences of strain XHU 5135 and other related species is shown in Fig. 5 with high levels of bootstrap support. In addition, *Aidingimonas halophila* YIM 90637 [40], *Streptomyces aidingensis* TRM46012 [42], *Longimycelium tulufanense* TRM 46004 [43], *Salinifilum aidingensis* TRM 46074 [44], and *Gracilibacillus aidingensis* YIM 98001 [45]were also isolated from Aiding Lake, and above these strains have been characterized as some novel species. These results indicate that there are potentially unique, novel sources of bacteria in Aiding Lake.

**Fig 2.**
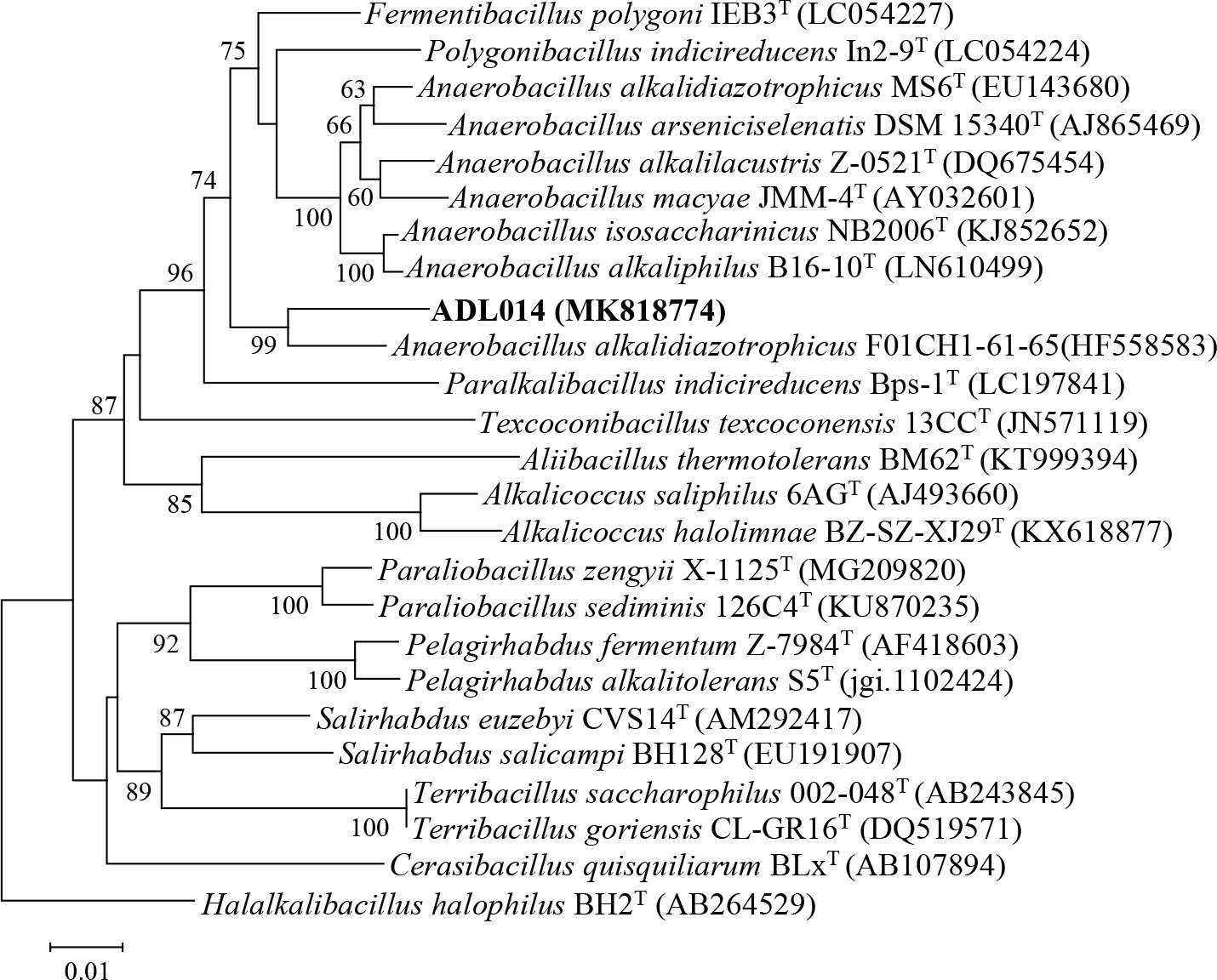
Phylogenetic tree of strain ADL014 and its near neighbors calculated from 16S rRNA gene sequences using Kimura’s evolutionary distance method (Kimura, 1980) and the neighbor-joining method of Saitou and Nei (1987). Bar, 0.01 nucleotide substitutions per site.

**Fig 3.**
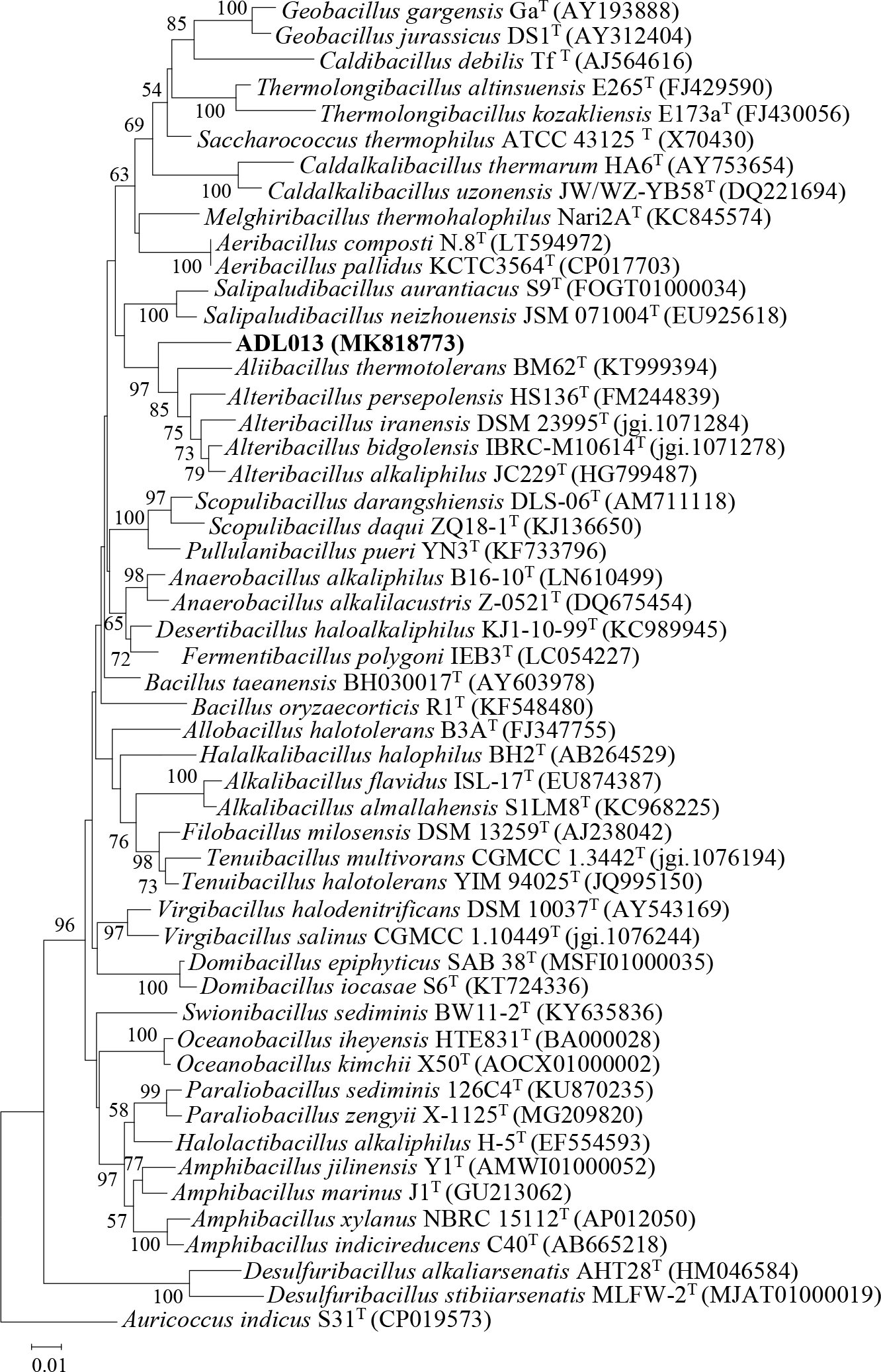
Phylogenetic tree of strain ADL013 and its near neighbors calculated from 16S rRNA gene sequences using Kimura’s evolutionary distance method (Kimura, 1980) and the neighbor-joining method of Saitou and Nei (1987). Bar, 0.01 nucleotide substitutions per site.

**Fig 4.**
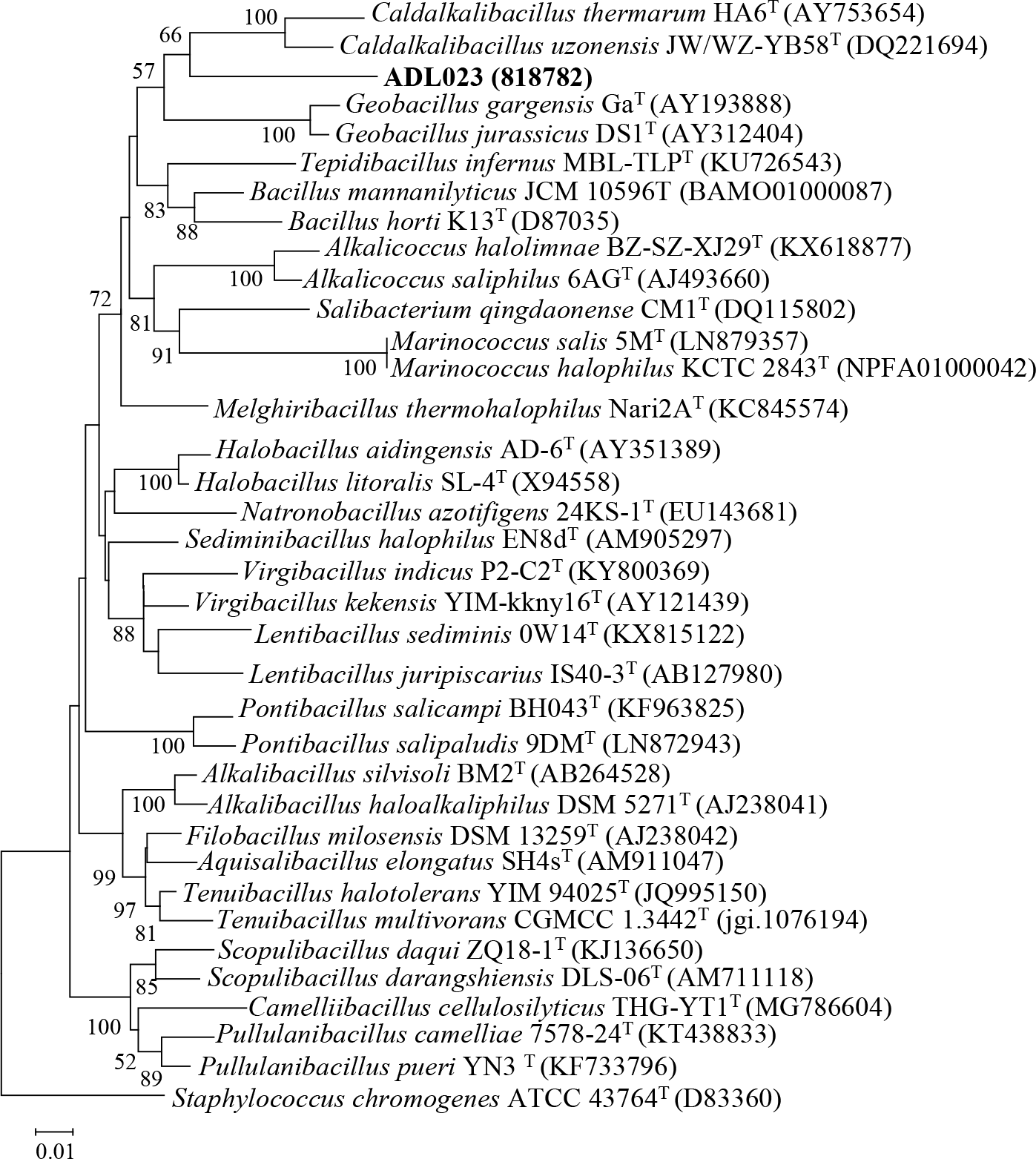
Phylogenetic dendrogram for taxa of the family Bacillaceae reconstructed using the neighbor-joining method based on almost complete 16S rRNA gene sequences to display the taxonomic position of strain ADL023. *Staphylococcus chromogenes* ATCC 43764^T^ (D83360) was used as the root organism. Numbers at nodes indicate levels of bootstrap support (%) based on neighbor-joining analysis of 1000 resampled datasets; only values above 50% are shown. Bar, 0.01 nucleotide substitutions per site.

**Fig 5.**
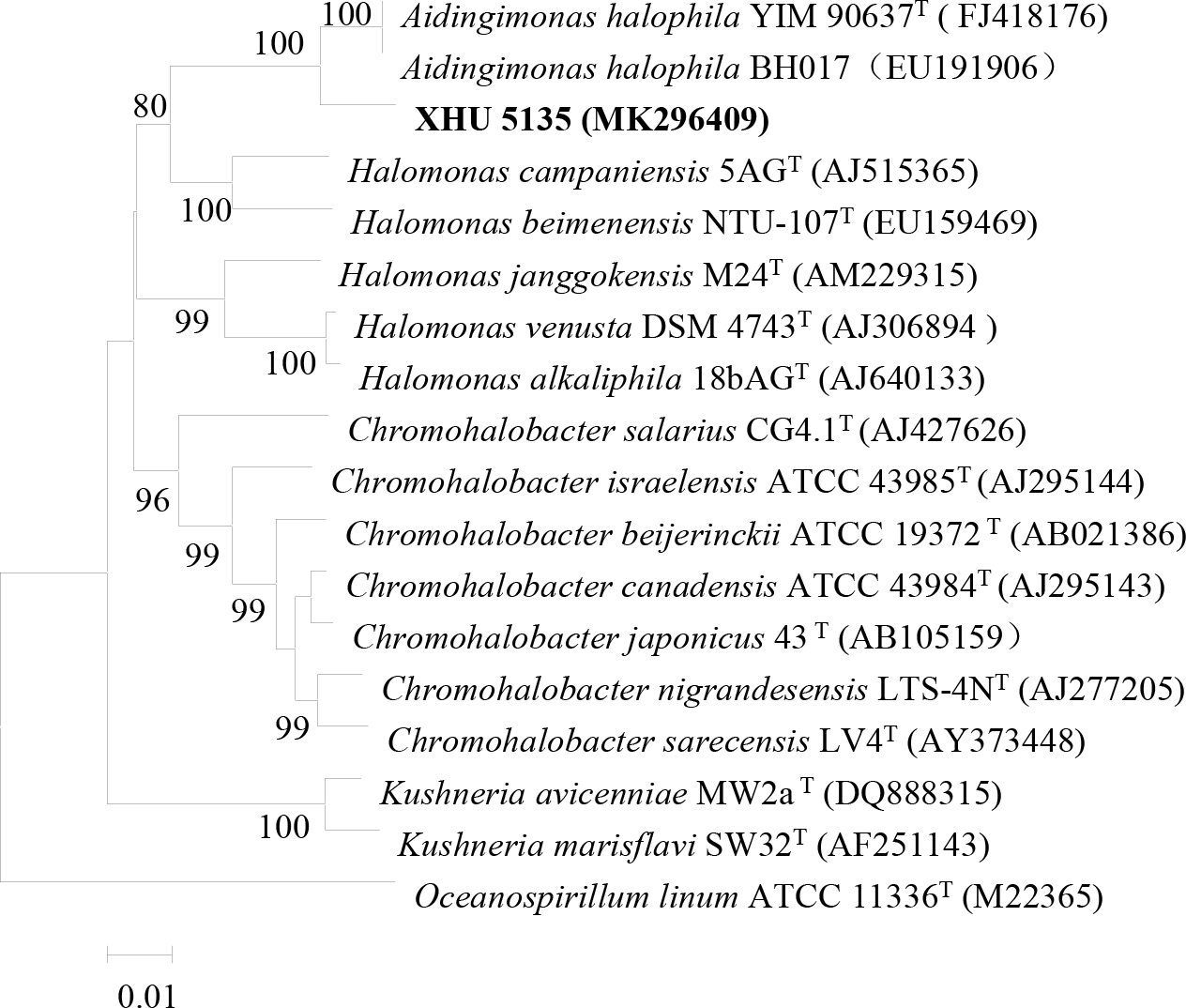
Neighbour-joining tree based on 16S rRNA gene sequences, showing the phylogenetic relationships of the novel isolate XHU 5135 and related taxa. Numbers at nodes are bootstrap percentages based on 1000 replicates. Bar, 0.01 nucleotide substitutions per site.

Generally, bacterial diversity in extreme environments is relatively low. However, in the study, an unexpectedly high bacterial diversity was observed using sediment samples from Aiding Lake. At least 41 bacterial genera were identified, suggests that some strains might represent a valuable source of new species, thereby providing a new reference for further understanding bacterial diversity in hypersaline environments. This study also demonstrated that the diversity of bacteria isolated from Aiding Lake is largely dependent on the isolation media. No universal medium or uniform isolation technology has been established for microbial resources around the world. Therefore, there is a need to develop novel isolation media or isolation techniques to better mine non-culturable bacterial resources.

## Acknowledgments

The National Natural Science Foundation of China (Project No. 30660005) and Office of education in Sichuan Province, China (Project Nos. 13205688, 13ZB0024) supported this study.

## References

1. Oren A. Industrial and environmental applications of halophilic microorganisms. Environ Technol 2010; 31: 825–834. https://doi.org/10.1080/09593330903370026

2. Oren A. Halophilic microbial communities and their environments. Curr Opin Biotech 2015; 33: 119–124. https://doi.org/10.1016/j.copbio.2015.02.005

3. Oren A. Molecular ecology of extremely halophilic Archaea and Bacteria. FEMS Microbiol Ecol 2003; 39(1): 1–7. https://doi.org/10.1111/j.1574-6941.2002.tb00900.x

4. Antonio V, Rafael RLH, Cristina SP, Thane PR. Microbial diversity of hypersaline environments: a metagenomic approach. Curr Opin Microbiol 2015; 25: 80–87. https://doi.org/10.1016/j.mib.2015.05.002

5. Ali N, Giti E, Mohammad AA, Mariana SC, Lucas JS, Zahra E, Seyed ASF, et al. Microbial diversity in the hypersaline lake meyghan, Iran. Scientific Reports 2017; 7: 11522. https://doi.org/10.1038/s41598-017-11585-3

6. Manel BA, Fatma K, Najwa K, Fabrice A, Najla M, Marianne Q, et al. Abundance and diversity of prokaryotes in ephemeral hypersaline lake Chott El Jerid using Illumina Miseq sequencing, DGGE and qPCR assays. Extremophiles 2018; 22(5): 811–823. https://doi.org/10.1007/s00792-018-1040-9

7. Nahid O, David JC, Inmaculada L, Victoria B, Fernando MC. Study of bacterial community composition and correlation of environmental variables in Rambla Salada, a hypersaline environment in south-eastern Spain. Front Microbiol 2018; 9: 1377–1383. https://doi.org/10.3389/fmicb.2018.01377

8. DasSsrma S, Arora P. 2001. Halophiles, p. 458–466. In J. R. Battista et al. (ed.), Encyclopedia of life sciences, vol. 8. Nature Publishing Group, London, Untied Kingdom.

9. Oren A. The bioenergetics basis for the decrease in metabolic diversity ecosystems. Hydrobiologia 2001; 466: 61–72.

10. Meglio LD, Santos F, Gomariz M, Almansa C, Lopez C, Anton J, et al. Seasonal dynamics of extremely halophilic microbial communities in three argentinian salterns. FEMS Microbiol Ecol 2016; 92(12): 184. https://doi.org/10.1093/femsec/fiw184

11. Oren A. Diversiyt of halophilic microorganisms: environments, phylogeny, physiology, and applications. J Ind Microbiol Biotechnol 2002; 28(1): 56–63. https://doi.org/10.1038/sj/jim/7000176

12. Addis S, Anders L, Amare G, Lise O. Prokaryotic community diversity along an increasing salt gradient in a soda Ash concentration Pond. Microbiol Ecology 2016; 71(2): 326–338. https://doi.org/10.1007/s00248-015-0675-7

13. Jiang HC, Dong HL, Zhang GX, Yu BS, Leah RC, Matthew WF. Microbial diversity in water and sediment of lake Chaka, an athalassohaline lake in northwestern China. Appl Environ Microbiol 2006; 72(6): 3832–3845. https://doi.org/10.1128/AEM.02869-05

14. Loubna T, Donald PB, Alan RH, Keith AC. Life in extreme environments: microbial diversity in Great Salt Lake, Utah. Extremophiles 2014; 18(3): 525–535. https://doi.org/10.1007/s00792-014-0637-x

15. Han R, Zhang X, Liu J, Long QF, Chen LS, Liu DL, et al. Microbial community structure and diversity within hypersaline Keke salt lake environments. Can J Microbiol 2017; 63(11): 895–908. https://doi.org/10.1139/cjm-2016-0773

16. Baxter BK. Great Salt Lake microbiology: a historical perspective. Int Mcrobiology 2018; 21(3): 79–95. https://doi.org/10.1007/s10123-018-0008-z

17. Tang SK, Wang Y, Guan TW, Lee JC, Kim CJ, Li WJ. *Amycolatopsis halophila* sp. nov., a novel halophilic actinomycete isolated from a salt lake in China. Int J Syst Evol Microbiol 2010; 60(5): 1073–1078. https://doi.org/10.1099/ijs.0.012427-0

18. Guan TW, Zhao K, Xiao J, Liu Y, Xia ZF, Zhang XP, et al. *Brevibacterium salitolerans* sp. nov., an actinobacterium isolated from a salt lake in Xinjiang, China. Int J Syst Evol Microbiol 2010; 60: 2991–2995. https://doi.org/10.1099/ijs.0.020214-0

19. Guan TW, Wei B, Zhang Y, Xia ZF, Che ZM, Chen XG, et al. *Actinopolyspora lacussalsi* sp. nov., *Actinopolyspora lacussalsi* sp. nov., an exteremely halophilic actinomycete isolated from a salt lake. Int J Syst Evol Microbiol 2013; 63: 3009–3013. https://doi.org/10.1099/ijs.0.047167-0

20. Duangmal K, Suksaard P, Pathom-aree W, Mingma R, Matsumoto A, Takahashi Y, et al. *Actinopolyspora salinaria* sp. nov., a halophilic actinomycete isolated from solar saltern soil. Int J Syst Evol Microbiol 2016; 68(4): 3506–3511. https://doi.org/10.1099/ijsem.0.000926

21. Harjodh S, Manpreet K, Shweta S, Sunita M, Shekhar K, Venkata RV, et al. *Salibacterium nitratireducens* sp nov., a haloalkalitolerant bacterium isolated from a water sample from Sambhar salt lake, India. Int J Syst Evol Microbiol 2018; 68: 3506–3511. https://doi.org/10.1099/ijsem.0.003021

22. Meng XL, Ming H, Huang JR, Zhang LY, Cheng LJ, Zhao ZL, et al. *Paracoccus halotolerans* sp. nov., isolated from a salt lake. Int J Syst Evol Microbiol 2019: 69: 523–528. https://doi.org/10.1099/ijsem.0.003190

23. Yakimov M, Giuliano L, Crisafi E, Chernikova T, Timmis K, Golyshin P. Microbial community of a saline mud volcano at San Biagio-Belpasso, Mt. Etna (Italy). Environ Microbiol 2002; 4(5): 249–256. https://doi.org/10.1046/j.1462-2920.2002.00293.x

24. Shirling EB, Gottlieb D. Methods for characterization of Streptomyces species. Int J Syst Bacteriol 1966; 16: 313–340. https://doi.org/10.1099/00207713-16-3-313

25. Yoon SH, Ha SM, Kwon S, Lim J, Kim Y, Seo H, et al. Introducing EzBioCloud: A taxonomically united database of 16S rRNA and whole genome assemblies. Int. J. Syst. Evol. Microbiol 2017; 67: 1613–1617. https://doi.org/10.1099/ijsem.0.001755

26. Li WJ, Xu P, Schumann P, Zhang YQ, Pukall R, Xu LH, et al. *Georgenia ruanii* sp. nov., a novel actinobacterium isolated from forest soil in Yunnan (China), and emended description of the genus *Georgenia*. Int J Syst Evol Microbiol 2007; 57: 1424–1428. https://doi.org/10.1099/ijs.0.64749-0

27. Tamura K, Stecher G, Peterson, D, Filipski A, Kumar S. MEGA6: Molecular Evolutionary enetics Analysis Version 6.0. Mol Biol Evol 2013; 30(12): 2725–2729. https://doi.org/10.1093/molbev/mst197

28. Saitou N, Nei M. The neighbor-joining method: a new method for reconstructing phylogenetic tree. Mol Biol Evol 1987; 4(4): 406–425. https://doi.org/10.1093/oxfordjournals.molbev.a040454

29. Kimura M. A simple method for estimating evolutionary rates of base substitutions through comparative studies of nucleotide sequence. J Mol Evol 1980; 16(2): 111–120. https://doi.org/10.1007/BF01731581

30. Felsenstein J. Confidence limits on phylogenies: an approach using the bootstrap. Evolution 1985; 39: 783–789.

31. Ezaki T, Hashimoto Y, Yabuuchi E. Fluorometric deoxyribonucleic acid-deoxyribonucleic acid hybridization in microdilution wells as an alternative to membrane filter hybridization in which radioisotopes are used to determine genetic relatedness among bacterial strains. Int J Syst Bacteriol 1989; 39: 224–229.

32. Phillips K, Frederic 3rd Z, Elizondo OR, Lowe KL. Phenotypic characterization and 16S rDNA identification of culturable non-obligate halophilic bacterial communities from a hypersaline lake, La Sal del Rey, in extreme South Texas (USA). Aquatic. Biosystems 2012; 8(1): 5–15. https://doi.org/10.1186/2046-9063-8-5

33. Li X, Yu YH. Biodiversity and screening of halophilic bacteria with hydrolytic and antimicrobial activities from yuncheng salt lake, China. Biologia 2015; 70(2): 151–156. https://doi.org/10.1515/biolog-2015-0033

34. Rohban R, Amoozegar MA, Ventosa A. Screening and isolation of halophilic bacteria producing extracellular hydrolyses from Howz Soltan Lake, Iran. J Ind Microbiol Biot 2009; 36(3): 333–340. https://doi.org/10.1007/s10295-008-0500-0

35. Ellis GA, Thomas CS, Chanana S, Adnani N, Szachowicz E, Braun DR, et al. Brackish habitat dictates cultivable actinobacterial diversity from marine sponges. Plos One 2017; 12(7): e0176968. https://doi.org/10.1371/journal.pone.0176968

36. Zhang GY, Cao TF, Ying JX, Yang YL, Ma LQ. Diversity and novelty of actinobacteria in Arctic marine sediments. Antonie. Van. Leeuwenhoek 2014; 105(4): 743–754. https://doi.org/10.1007/s10482-014-0130-7

37. Atika M, Nasserdine S, Abdelghani Z, Florence M, Ahmed L. Isolation, taxonomy, and antagonistic properties of halophilic actinomycetes in Saharan soils of Algeria. Appl Environ Microbiol 2011; 77(18): 6710–6714. https://doi.org/10.1128/AEM.00326-11

38. Ronoh RC, Budambula NLM, Mwirichia RK, Boga HI. Isolation and characterization of actinobacteria from lake Magadi, Kenya. Afr J Microbiol Res 2013; 7: 4200–4206. https://doi.org/10.5897/AJMR2012.2296

39. Stackebrandt E, Goebel BM. Taxonomic note: a place for DNA-DNA reassociation and 16S rRNA sequence analysis in the present species definition in bacteriology. Int J Syst Bacteriol 1994; 44: 846–849. https://doi.org/10.1099/00207713-44-4-846

40. Wang Y, Tang SK, Lou K, Lee JC, Jeon CO, Xu LH, et al. *Aidingimonas halophila* gen. nov., sp. nov., a moderately halophilic bacterium isolated from a salt lake. Int J Syst Evol Microbiol 2009; 59(12): 3088–3094. https://doi.org/10.1099/ijs.0.010264-0

41. Wayne LG, Brenner DJ, Colwell RR, et al. International Committee on Systematic Bacteriology. Report of the ad hoc committee on reconciliation of approaches to bacterial systematics. Int J Syst Bacteriol 1987; 37: 463–464.

42. Xia ZF, Ruan JS, Huang Y, Zhang LL. *Streptomyces aidingensis* sp. nov., an actinomycete isolated from lake sediment. Int J Syst Evol Microbiol 2013; 63(9): 3204–3208. https://doi.org/10.1099/ijs.0.049205-0

43. Xia ZF, Guan TW, Ruan JS, Huang Y, Zhang LL. *Longimycelium tulufanense* gen. nov., sp. nov., a filamentous actinomycete of the family Pseudonocardiaceae. Int J Syst Evol Microbiol 2013; 63(8): 2813–2818. https://doi.org/2813-2818.10.1099/ijs.0.044222-0

44. Nikou MM, Ramezani M, Harirchi S, Makzoom S, Amoozegar MA, Shahzadeh FSA, et al. Salinifilum gen. nov., with description of *Salinifilum proteinilyticum* sp nov., an extremely halophilic actinomycete isolated from Meighan wetland, Iran, and reclassification of *Saccharopolyspora aidingensis* as *Salinifilum aidingensis* comb. nov and *Saccharopolyspora ghardaiensis* as *Salinifilum ghardaiensis* comb. Nov Int J Syst Evol Microbiol 2017; 67(10): 4221–4227. https://doi.org/10.1099/ijsem.0.002286

45. Guan TW, Tian L, Li EY, Tang SK, Zhang XP. *Gracilibacillus aidingensis* sp. nov., a novel moderately halophilic bacterium isolated from Aiding salt lake. Arch Microbiol 2017; 199(9): 1277–1281. https://doi.org/10.1007/s00203-017-1399-5

